# Comparable Strength and Hypertrophic Adaptations to Low-Load and High-Load Resistance Exercise Training in Trained Individuals: Many Roads Lead to Rome

**DOI:** 10.1101/2025.04.28.650925

**Authors:** Kristoffer Toldnes Cumming, Ingrid Cecelia Elvatun, Richard Kalenius, Gordan Divljak, Truls Raastad, Niklas Psilander, Oscar Horwath

**Author notes:** joint senior authors (Niklas Psilander, Oscar Horwath).

## Abstract

The muscular and myocellular adaptations to low-load resistance exercise training (LL-RET) remain incompletely understood, particularly in the trained state. The primary aim of this study was to examine adaptations to an LL-RET regimen and compare these to a high-load training regimen (HL-RET).

Fourteen resistance-trained males and females (26.4 ± 4.4 years) participated in a 9-week RET program (twice per week). Using a within-subject design, each individual trained one leg with HL-RET (3-5 repetitions), and the other with LL-RET (20-25 repetitions), all sets performed to volitional failure.

Pre- and post-intervention, muscle endurance, maximal strength, and muscle thickness using ultrasound was assessed. Muscle biopsies were analyzed for fiber type composition, fiber cross-sectional area (fCSA), and satellite cell- and myonuclear content using immunofluorescence.

The training regimens led to comparable increases in maximal strength in multi-joint movements (21%), but not in single-joint movements were HL-RET was superior. LL-RET induced superior improvements in local muscle endurance (9% vs -2.7%, p=0.013). Regardless of training regimen, muscle thickness increased by ∼7.4% at the mid-thigh site and ∼8.5% at the distal site pre-to post-intervention. However, no changes were observed in fiber type composition or fCSA. Satellite cell content increased by ∼25% in type I fibers, independent of training regimen, but no changes were noted in myonuclear content.

Here we novelly show that LL-RET can replicate many aspects of HL-RET leading to similar increases in both muscle hypertrophy and strength. Our study thus supports the notion that comparable adaptations to RET can be achieved using distinct loading regimens.

**New and noteworthy:** This study compared two distinct resistance exercise loading strategies (3-5 RM vs. 20-25 RM) in trained individuals, evaluating both muscular and myocellular adaptations. Our findings demonstrate that low-load resistance exercise training (LL-RET) is an effective alternative to traditional high-load strategies for increasing strength and muscle size. These results highlight that skeletal muscle growth can be achieved through various external stressors, offering valuable insights for individuals seeking hypertrophy but unable to tolerate high loads.

## Introduction

Skeletal muscle mass accounts for ∼30-40% of total body weight in healthy adults, making it the largest component of the human body. In addition to its fundamental role in locomotion, skeletal muscle is a key regulator of metabolic homeostasis, nutrient disposal, and having more muscle mass is beneficial for the recovery from illness or trauma (20, 30, 51). Consequently, promoting growth and maintenance of muscle mass is essential for supporting an active lifestyle and reducing the risk of metabolic disorders across the entire lifespan (58). Conversely, muscle loss is associated with frailty, reduced capacity for daily activities, and an increased risk of metabolic diseases, including obesity and diabetes (27, 50, 53).

Resistance exercise training (RET) is widely accepted as the most efficient non-pharmacological intervention for stimulating muscle growth and enhancing muscular strength (17). Traditionally, it has been suggested that loads exceeding 70% of one repetition maximum (1 RM) are necessary to optimally induce adaptations to RET, i.e., strength gains and muscle growth (2). This is based on the principle of motor unit recruitment, where high-threshold motor units are activated only with loads ≥70% of 1 RM (18). However, this paradigm may not be applicable to some clinical populations, such as older adults with arthritis or athletes recovering from orthopedic surgery, who are unable to tolerate the mechanical stress associated with high-load RET (HL-RET). Given the limitations of traditional exercise prescriptions, alternative RET strategies merit further investigation. This is crucial not only for optimizing athletic performance and development but also for enhancing general health.

In recent years, low-load resistance exercise training (LL-RET) has emerged as a viable alternative to traditional high-load regimens (15, 19, 33, 37, 48, 54). However, it remains unclear whether LL-RET induces strength and hypertrophic adaptations comparable to those achieved with HL-RET, as findings across studies have been inconsistent (10, 19, 22, 38, 48). The reasons for these discrepancies are not fully understood but may be attributed to variations in study design, such as differences in training status, the total volume of weight lifted, and whether exercises are performed to momentary muscle fatigue (failure) or not. For example, Campos *et al*. (2002) found that moderate-and high-load conditions (3-11 RM) were more effective than low-load conditions (20-28 RM) in promoting strength and muscle fiber hypertrophy in untrained individuals, provided that training volume was equated across conditions (10). However, when training volume was not equated, i.e., the total volume was higher in the low-load condition, similar hypertrophic responses were observed in untrained subjects performing LL-RET (30% of 1 RM) and HL-RET (80% of 1 RM) (35). Morton *et al*. (2016) reported that low-load (20-25 RM) and moderate-load (8-12 RM) training performed to volitional failure produced similar strength gains and muscle fiber growth in resistance-trained individuals (38). Nevertheless, as Morton and colleagues did not include a high-load condition, i.e., 3-5 RM, it remains to be determined whether LL-RET can fully replicate the strength and hypertrophic response observed with HL-RET in trained populations when sets are taken to volitional failure (38). In addition, it is yet to be determined whether LL-RET induces a distinct hypertrophic response in terms of muscle fiber types. Some evidence implies that low-load regimens may preferentially stress type I fibers (4, 11), but a clear consensus remains to be established (19, 47).

The addition of myonuclei through satellite cell fusion serves as a key process driving muscle growth across various experimental models (8, 28, 49). In humans, the rate of myonuclear accretion is thought to dictate the rate of muscle growth in response to a period of HL-RET (44). However, much of our current understanding of the mechanisms underlying muscle growth in LL-RET stems from studies that combine LL-RET with blood-flow restriction (BFR). This combination effectively stimulates satellite cell proliferation and myonuclear accretion, particularly in type I fibers (4-6, 14, 26, 43). Yet, due to limited research on LL-RET alone, it remains unclear if this mechanism is central to muscle growth induced by this type of training. One study found that LL-RET alone, performed at 30% of 1 RM, induced robust increases in satellite cell numbers 48 hours after a single exercise session (55). However, it remains uncertain whether this temporary increase in satellite cell number leads to myonuclear accretion over time and whether these effects differ from those of HL-RET.

Therefore, using a within-subject design, the primary aim of this study was to assess the muscular and myocellular effects of a 9-week LL-RET intervention in resistance-trained individuals and compare it to HL-RET when training was performed to volitional failure. The secondary aim of this study was to determine potential fiber type-specific adaptations to the two different training protocols. We hypothesized that the magnitude of muscular adaptations would be similar between conditions, but that HL-RET would induce greater increases in muscle strength compared to LL-RET. We also hypothesized that LL-RET would stimulate adaptations primarily in type I fibers, whereas HL-RET would preferentially stimulate type II fibers.

## Methods

### Ethical approval

Before participating in the study, all participants were thoroughly informed about the nature of experimental procedures and the potential benefits and risks involved. Written informed consent was obtained from each participant before participating in the study. The study adhered to the ethical principles outlined in the Declaration of Helsinki and received approval from the local ethics committee (The Swedish Ethical Review Authority, DNR 2016/2159-31).

### Participants

The participants were recruited through social media, posters, and flyers at the Swedish School of Sport and Health Sciences in Stockholm, Sweden. Eligibility criteria included being between 20 and 35 years of age and free of diseases or musculoskeletal injuries to the lower extremities, as determined by a detailed medical history questionnaire. Participants were also required to have a history of strength training of at least 2 years, with the primary goal of increasing muscle mass and strength. This included performing at least one session per week targeting the lower body. Exclusion criteria included smoking or the use of medications that could affect muscular adaptations. Sixteen individuals initially volunteered for the study; however, two dropped out due to injuries sustained outside the intervention. Thus, 11 males and 3 females (n=14) completed the study. The participant’s physical characteristics are provided in Table 1.

**Table 1.**
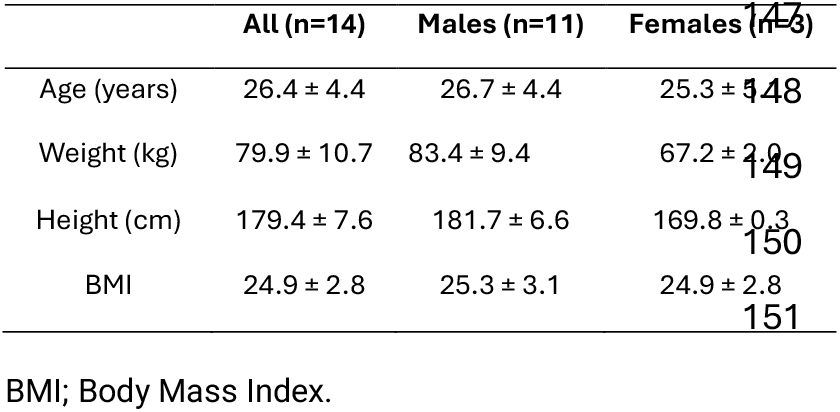
Physical characteristics of the participants.

### Experimental design

A schematic overview of the experimental design can be seen in Figure 1. This study used a within-subject design, where the participant’s legs were randomly assigned to one out of two unilateral training protocols: high-load (HL-RET) or low-load (LL-RET) resistance exercise training. A detailed description of the two training protocols is provided below. The unilateral design has been used in studies with similar research questions (22, 35) and allows us to control for training status, diet, and sleep, which all influence training adaptability (32). In total, the entire experimental period lasted for 11 weeks, where the first week (baseline) and the last week (re-tests) involved tests of muscle strength, muscle thickness, and muscle biopsy sampling. In turn, the training period lasted 9 weeks and was divided into two 4-week blocks interspersed by a one-week deload period with reduced training volume, see Figure 1. The deload period was added to the training program to reduce general fatigue associated with high-intensity training and to facilitate long-term muscle adaptability (3).

**Figure 1.**
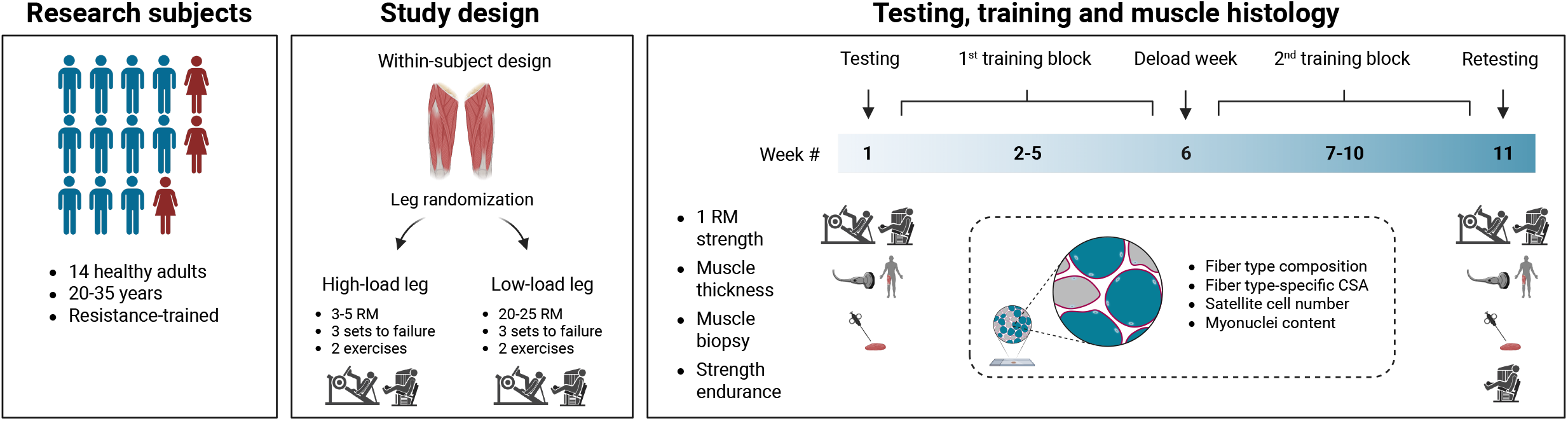
Schematic overview of the study design. This figure was created using BioRender.com.

### Training protocol

The training protocol consisted of two weekly sessions scheduled on Mondays and Thursdays. The exercises performed included leg press and leg extension. Each session began with a general warm-up on a stationary bicycle at a self-selected intensity for 5 minutes, followed by a specific warm-up on the leg press using standardized procedures (29). This warm-up was only performed for the leg subjected to the high load, as no warm-up was considered necessary for the LL-RET leg. After the warm-up, the HL-RET leg performed 3-5 repetitions, corresponding to ∼90-95% of the individual 1 RM, while the LL-RET leg performed 20-25 repetitions, corresponding to ∼40-60% of the individual 1 RM. Each set was taken to volitional failure, accompanied by strong verbal encouragement. Each leg performed 3 sets per exercise, with a 2-minute rest between sets. Within the session, the legs were trained alternately, e.g., one set of HL-RET was followed by one set of LL-RET. If the repetitions deviated from the prescribed range during an exercise, the load was adjusted for the subsequent set. During the intervention, the load was adjusted according to reflect the targeted percentage of 1 RM. The leg that was selected to start the exercise session was randomly selected at the start of the intervention and alternated every week thereafter. During the deload week, only one exercise session was conducted. In this session, the number of sets per leg and exercise was reduced to one, but the intensity and repetition ranges were kept constant.

All repetitions were performed in a controlled manner using the full range of motion. The leg press exercise was correctly performed when the sled was eccentrically lowered until it reached a 90° knee angle and concentrically pushed up until full extension. The leg extension exercise was correctly performed at a minimum of 160° knee angle in the top position. In total, the training protocol consisted of 17 sessions, all supervised by experienced trainers and investigation team members. The average training volume for each leg was calculated by multiplying the number of repetitions with the load lifted during each session. This was later summed up and divided by the number of sessions attended, ultimately expressed in kgs, as reported elsewhere (38).

### Training intervention period

During the training period, the participants were instructed to refrain from all strength training-related activities for the lower body but were allowed to perform strength-and endurance training for the upper body. To ensure adequate protein intake throughout the intervention, the participants immediately consumed one serving of high-quality whey protein after each training session (137 kcal, 27 g protein, 2.5 g carbohydrates, 2.1 g fat, Tyngre© AB, Johanneshov, Sweden). The participants were also instructed to maintain their dietary habits throughout the intervention.

### Testing procedures

One week before the start and one week after the intervention period ended, the participants underwent a test battery consisting of 1 RM strength testing, local muscle endurance (only after the intervention), measures of muscle thickness using ultrasound, and muscle biopsies sampled from the *vastus lateralis* muscle, see Figure 1. Specific information on each of these procedures is provided in the sections below. In addition, the participants were instructed to refrain from nicotine and nutritional supplements for at least three hours before testing and to avoid vigorous activities for a minimum of 48 hours before testing.

### Anthropometric measures

The participant’s height (cm) and weight (kg) were measured using standardized procedures with a calibrated stadiometer and digital scale, respectively.

### Maximum strength testing

One repetition maximum (1 RM) strength testing was conducted in the inclined leg press and the leg extension, both sets of equipment purchased from Cybex, Träningsredskap AB, Jönköping, Sweden. After a brief general warm-up, a specific warm-up of the exercise was performed at ∼50% of the participant’s estimated 1 RM based on the 10 RM testing. The weight was then progressively increased by 10% after each successful attempt, continuing until the participant was unable to lift the weight with proper technique (29). A three-minute rest period was provided between each attempt. In the leg press exercise, the participant had to lower the sled to a 90° angle at the knee joint and then reach full extension to complete one repetition. In the leg extension exercise, the participant had to lower the weight to the bottom position and extend the lower leg to a minimum of 160° to complete one repetition. The 1 RM test was performed for each leg individually.

### Strength endurance testing

After the study, a local muscular endurance test was conducted on the leg extension exercise. Due to the associated metabolic stress, this test was performed after the 1 RM strength assessment. Both the HL-RET and the LL-RET legs were required to perform as many repetitions as possible with the load used during the final session of the LL-RET program. Each leg completed a single set, with a 2-minute rest interval between the legs. Participants were instructed to continue until muscular failure while maintaining proper form. The test was terminated if the subject failed to fully extend the leg between 160° and 180° in consecutive repetitions. The starting leg for each test was randomly selected.

### Muscle thickness

Muscle thickness was measured using a real-time two-dimensional (2D) B-mode ultrasound (Philips Ultrasound, Andover, Massachusetts, USA) with a 40 mm wide probe operating at 11 Hz. Muscle thickness was defined as the distance between the deep and superficial aponeurosis. First, with the ultrasound probe placed in the transversal plane and perpendicular to the skin, the muscle thickness of the muscle belly of *vastus lateralis* was measured at 50% of the distance between the lateral epicondyle of the femur and the distal portion of the trochanter major (MID). Second, muscle thickness of the same muscle was measured at 22% of the distance from the lateral epicondyle of the femur to the distal portion of the trochanter major (DIST). The measurement site was carefully determined using a measuring tape and thereafter highlighted with a permanent marker to ensure that subsequent measures could be obtained from the same anatomical site at the post test. Minimal pressure was applied to the probe to avoid changing the tissue architecture. One image was saved for each anatomical measurement site, and images were later analyzed using ImageJ (National Institutes of Health, Bethesda, MD, USA). An average of three measurements per image was obtained from each site. All measurements were performed in a single-blind manner by an experienced operator.

### Muscle sampling

Muscle tissue was sampled from all participants at least 2 days before baseline testing and at least 2 days after retesting at week 11. After applying local anesthesia (Carbocain, 20 mg . ml^-1^, AstraZeneca, Södertälje, Sweden), a small incision was made to the skin and fascia, after which muscle samples were obtained from the midpart of the *vastus lateralis* muscle using the Weil-Blakesley conchotome technique (16). After excision, the muscle pieces were rapidly cleaned free of excessive blood, connective-and adipose tissue. The samples were then embedded in OCT (OCT Cryomount, Histolab, Sweden) and frozen in isopentane cooled by liquid nitrogen. The samples were stored in a freezer (−80°C) until sectioning commenced. The pre-post incisions were separated laterally by at least ∼3 cm.

### Immunofluorescence

Cross-sections (8 µm) were prepared from the OCT-embedded samples using a cryostat (CM1860 UV, Leica Microsystems, Nussloch, Germany) and carefully placed on microscope slides (SuperFrost Plus, Thermo Fisher, Braunschweig, Germany). The cross-sections were left to air-dry for 1h before being stored in the freezer (−80°C). All four samples collected from each individual were mounted together to minimize variability, i.e., pre-post samples from the HL and the LL leg.

The immunohistochemical procedures related to the staining of myosin heavy chain (MyHC), satellite cells, and myonuclei have been described in previous publications from our laboratory (12, 23, 45). Information on antibodies used in the present study is listed in Table 2. Briefly, slides to determine muscle fiber type composition and fiber cross-sectional area (fCSA) were blocked for 60 mins in 1% bovine serum albumin (BSA) in phosphate-buffered saline (PBS) containing 0.05% Tween-20. The slides were then incubated overnight (4°C) with a mixture of primary antibodies against MyHC I (1:500, BA-D5) purchased from Developmental Studies of Hybridoma Bank (DSHB) and dystrophin (1:500, Ab15277, Abcam), see Table 2. The next day, slides were washed in PBS and incubated for 60 mins at room temperature with species-specific secondary antibodies purchased from Invitrogen (Alexa Fluor anti-mouse IgG 488 and anti-rabbit IgG 594), see Table 2. After this step, the slides were re-washed in PBS, mounted with ProLong Gold Antifade medium containing DAPI (P36935, Molecular Probes), and covered with a coverslip.

**Table 2.** Information on antibodies used for immunofluorescence.

| Target | Primary | Species | Company | Subclass | Dilution | Secondary | Dilution |
| --- | --- | --- | --- | --- | --- | --- | --- |
| MyHC-I | BA-D5 | Mouse | DSHB | IgG2b | 1:500 | AF GAM 488 | 1:200 |
| Dystrophin | Ab15277 | Rabbit | Abcam | IgG | 1:500 | AF GAR 594 | 1:200 |
| Pax7 | Pax7 | Mouse | DSHB | IgG1 | 1:20 | AF GAM 488 | 1:200 |
| Laminin | Z0097 | Rabbit | DAKO | IgG | 1:500 | AF GAR 594 | 1:200 |
| PCM1 | HPA023370 | Rabbit | Sigma-Aldrich | IgG | 1:1000 | AF GAR 488 | 1:200 |
| Dystrophin | MANDYS8 | Mouse | DSHB | IgG2b | 1:20 | AF GAM 594 | 1:200 |

MyHC; Myosin Heavy Chain, DSHB; Developmental Studies of Hybridoma Bank, AF; AlexaFluor, GAM; Goat Anti-Mouse, GAR; Goat Anti-Rabbit, Pax7; Paired Box Protein 7, PCM1; Pericentriolar Material 1.

Slides to determine satellite cell content were first incubated with a solution containing 4% formaldehyde and 0.05 Triton-X-100 (Sigma Life Science, St. Louis, MO, USA) for 10 mins, washed in PBS, and blocked for 10 mins using a serum-free protein blocking solution (X0909, Dako, Glostrup, Denmark). The slides were then incubated overnight with primary antibodies against Pax7 (1:20, anti-Pax7, DSHB) and laminin (1:500, Z0097, Dako). The following day, slides were washed in PBS and incubated with secondary antibodies (Alexa Fluor anti-mouse IgG 488 and anti-rabbit IgG 594) diluted in 1% BSA in PBS. After this step, slides were washed in PBS, mounted in a DAPI-containing medium, and covered with a coverslip.

Slides to determine myonuclear content were first incubated for 30 mins with 2% BSA in PBS before being incubated overnight with primary antibodies against PCM1 (1:1000, HPA023370, Sigma Aldrich) and dystrophin (1:20, MANDYS8, DSHB) which were dissolved in 5% BSA and 0.02% of IGEPAL CA-630 (Sigma Aldrich, I3021, USA). The next day, slides were washed in PBS and then incubated at room temperature with secondary antibodies (Alexa Fluor anti-rabbit IgG 488 and anti-mouse IgG) diluted in 2% BSA. After the final washes, the slides were mounted with DAPI and covered with a coverslip. A representative image of this staining is provided in Figure 5G.

### Image acquisition and analyses

Stained sections were digitally captured using a microscope (BX61, Olympus, Tokyo, Japan) with a mounted camera (DP72, Olympus) using a 4× air objective (0.13 NA, UPlanFL N, Olympus) and a 10× air objective (0.30 NA, UPlanFL N, Olympus). Digital images were processed and analyzed for fiber type composition and size using the TEMA software (Checkvision, Hadsund, Denmark), and satellite cells and myonuclei were analyzed using ImageJ.

Muscle fiber type composition was determined from slides stained for MyHC-I and dystrophin, in which type I fibers were positive for the BA-D5 antibody, and type II fibers were left unstained, see Figure 4A. The average number of fibers included in this analysis per biopsy was 464 ± 172. The fCSA of type I and type II fibers were measured in fibers free from freezing artifacts. Fibers located at the periphery of the section were excluded from the analysis. This analysis contained an average of 83 ± 38 type I fibers and 86 ± 33 type II fibers per biopsy. Satellite cells refer to Pax7^+^/DAPI^+^ cells located inside the laminin border. The fiber type identity of the “parent” muscle fiber to which the satellite cell (and myonuclei) belongs was determined using serial sections stained for MyHC-I and dystrophin, as described above. To provide a reliable estimate of the satellite cell pool in human samples (34), we included an average of 497 ± 175 fibers per biopsy in this analysis. Moreover, to accurately quantify the myonuclear pool in these samples, we applied the specific myonuclear antibody PCM1 and double-stained nuclei together with the DAPI dye (57). As such, myonuclei refer to PCM1^+^/DAPI^+^ cells located within the border of the dystrophin staining. These cells were counted and related to their respective fiber type (as described earlier), including a minimum of 50 fibers per fiber type for each biopsy sample.

### Statistics

Data are presented as means ± standard deviation (SD) or means with individual values (pre-post). Baseline strength levels between the legs were compared using an unpaired t-test. The effects of the intervention were assessed using a two-way repeated measures analysis of variance (ANOVA) with factors for leg (LL-RET *vs* HL-RET) and time (pre *vs* post). Sidak’s multiple comparisons test was applied to localize the effects of each ANOVA model that revealed a significant interaction. Statistical analyses were conducted using GraphPad Prism (version 10.4.1 for Windows; GraphPad Software). Statistical significance was accepted at p ≤ 0.05. The study includes both males and females, but the sample size is too small for a sex-based comparison. However, data are presented separately by sex in the graphs to highlight potential differences and inform future research.

## Results

The compliance rate was 96%, with 228 out of 238 scheduled sessions completed for both legs. No participant missed more than two sessions. Ultrasound data were collected from all subjects (n=14). However, muscle biopsies from two subjects showed significant freezing artifacts and were excluded from the analysis (1M/1F). Accordingly, histological data is presented from 12 individuals (10M/2F).

### Training volume, 1 RM muscle strength and local muscle endurance

As planned, the average training volume per session was greater in the LL-RET compared to HL-RET, see Figure 2A. Specifically, the average training volume for the leg press exercise was 2.9-fold greater in LL-RET than in HL-RET (p<0.001). Similarly, the average training volume for the leg extension exercise was 1.4-fold greater in LL-RET compared to HL-RET (p<0.001).

**Figure 2.**
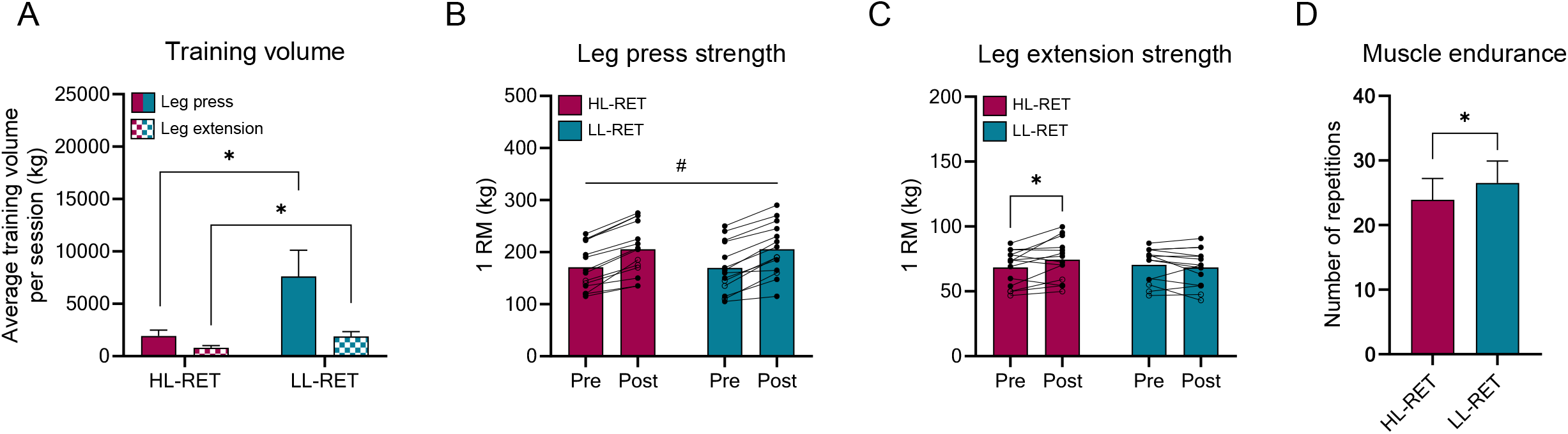
The average training volume for the HL-RET and LL-RET in the leg press exercise (solid colors) and the leg extension exercise (checkered colors) (A). This was calculated by multiplying the number of repetitions by the load lifted during each session. This was later summed up and divided by the number of sessions attended. 1 RM strength pre- and post-intervention in the leg press exercise (B) and the leg extension exercise (C). Local muscle endurance post-intervention in the HL-RET and the LL-RET (D.) Data are presented as means (bars) ± SD or with individual values (pre-post) from 14 subjects. Male subjects are presented as filled symbols (○), and female subjects are presented as unfilled (◯) symbols. Data was analyzed using two-way repeated measures ANOVA (A-C) and an unpaired t-test (D). Significantly different from HL-RET is indicated by * p<0.05 in A and D. Significantly different pre- to post-intervention is indicated by * p<0.05 in C. A significant main effect of time is indicated by # p<0.05 (B).

At baseline, the muscle strength (1 RM) in the leg press and the leg extension exercise did not differ between the conditions (HL-RET; leg press 171 ± 43 kg, leg extension 68 ± 14 kg, LL-RET; leg press 170 ± 48 kg, leg extension 70 ± 13 kg), see Figure 2B-C. Muscle strength (1 RM) in the leg press exercise increased on average by ∼21% from pre-to post-intervention (main effect of time; p<0.001) (Figure 2B). In the leg extension exercise, the HL-RET condition increased muscle strength (1 RM) from pre-to post-intervention by ∼9% (p=0.016), whereas no significant change was observed for the LL-RET condition (interaction effect; p=0.013, figure 2C). Local muscle endurance at post-intervention, determined as the number of repetitions performed until failure in the leg extension exercise, was greater in LL-RET than HL-RET (p=0.045), see Figure 2D.

### Muscle thickness

At baseline, ultrasound measures of muscle thickness did not differ between the legs/conditions in any of the two measuring sites (Figure 3A-3B). However, muscle thickness determined at the MID site increased on average by ∼7.4% from pre-to post-intervention (main effect of time; p=0.0198, figure 3A). Similarly, muscle thickness determined at the DIST site increased on average by ∼8.5% from pre-to post-intervention (main effect of time; p=0.025, figure 3B).

**Figure 3.**
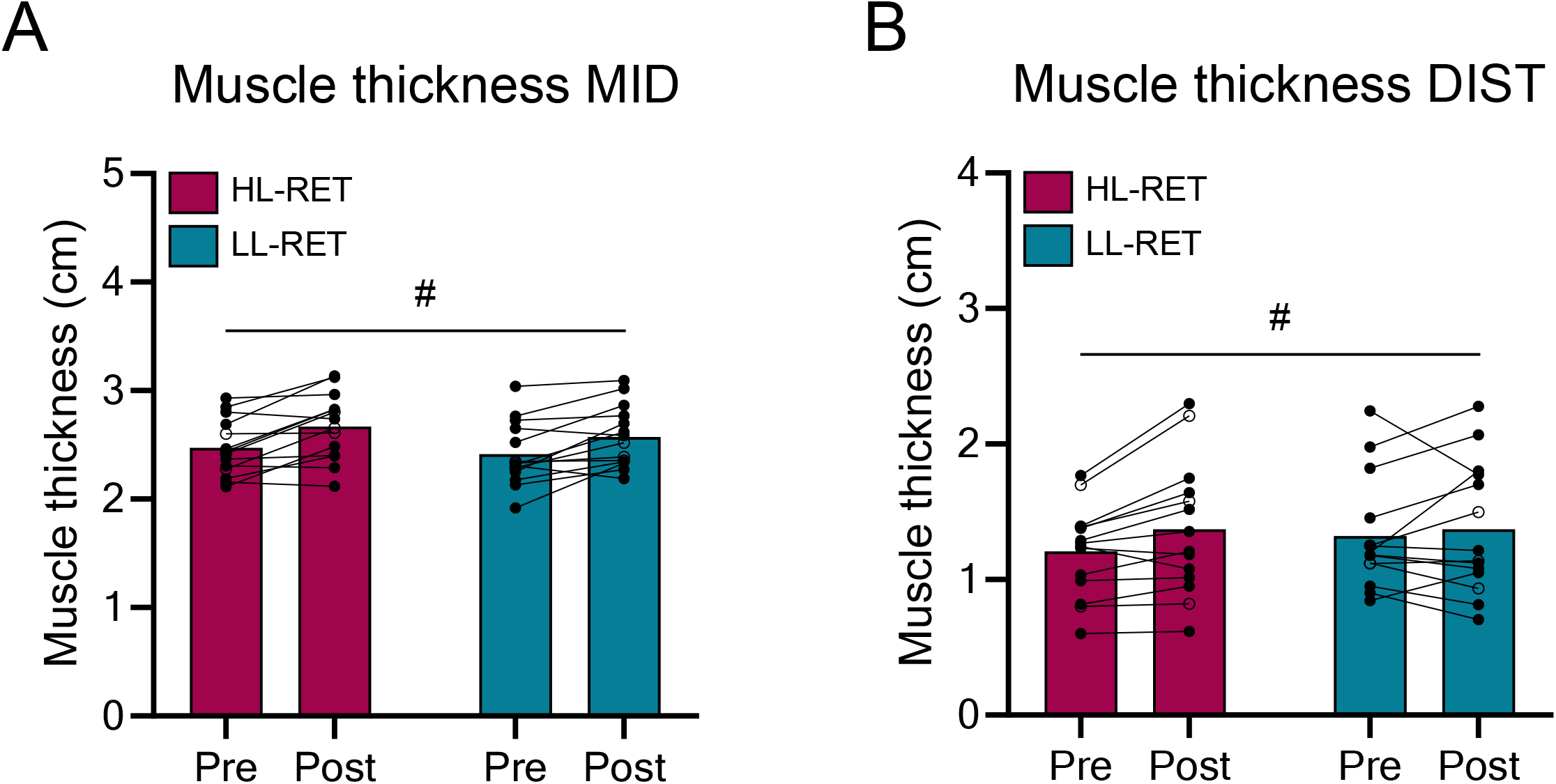
Muscle thickness pre- and post-intervention for HL-RET and LL-RET at the MID (A) and DIST (B) sites. Data are presented as means (bars) with individual values (pre-post) from 14 subjects. Male subjects are presented as filled symbols (○), and female subjects are presented as unfilled (◯) symbols. Data was analyzed using two-way repeated measures ANOVA. A significant main effect of time is indicated by # p<0.05.

### Fiber type composition and fCSA

The fiber type composition at baseline across both muscles was, on average, ∼42% and ∼58% type I and type II fibers, respectively, and this did not change from pre-to post-intervention, see Figure 4A. Similarly, no changes were observed for mixed, type I, and type II fCSA across the intervention period (Figure 4B, C, E). Additionally, an exploratory analysis using unpaired t-tests was conducted to compare delta changes in type I and type II fCSA between the two conditions, however, no significant differences were found (Figure 4D, F).

**Figure 4.**
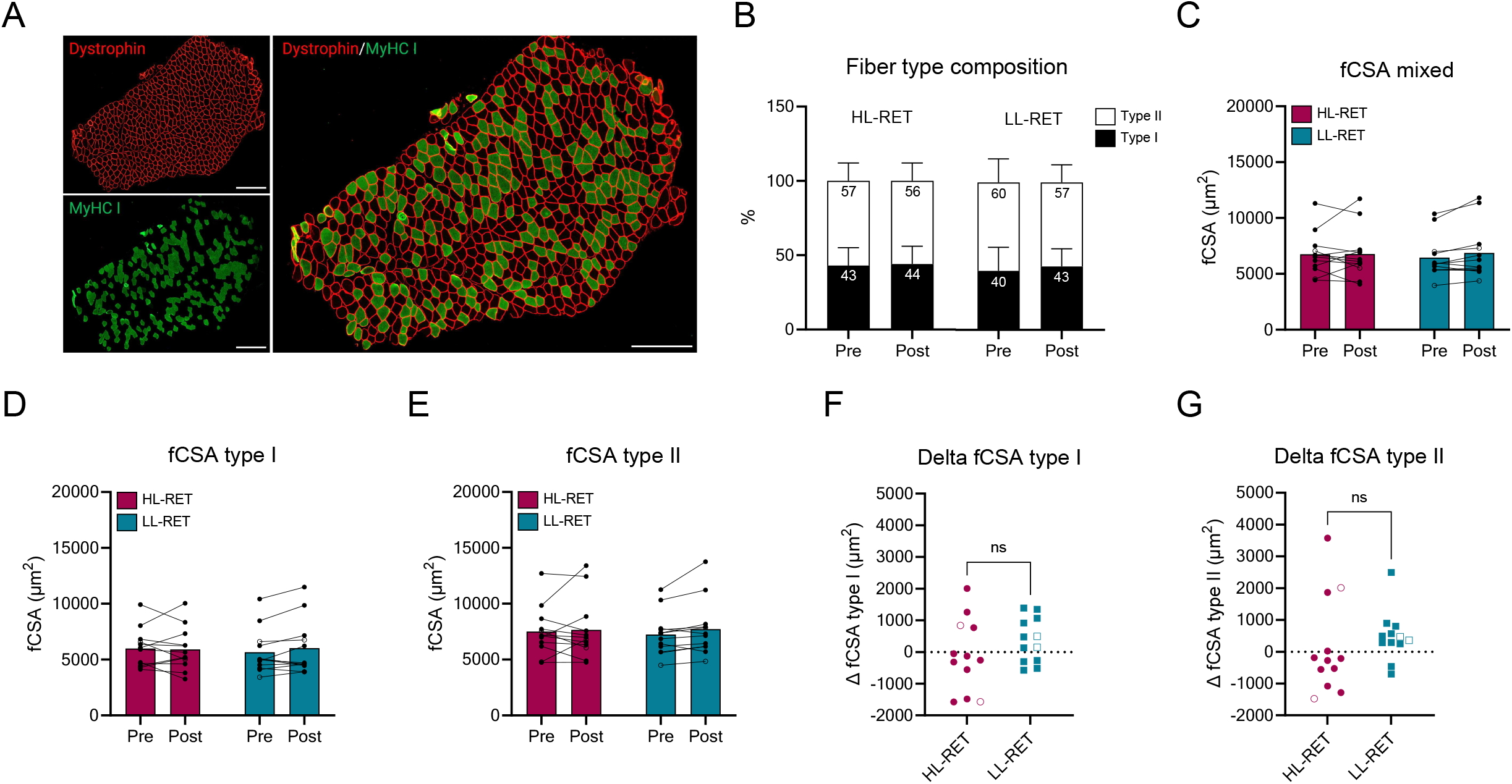
A representative image of the fiber type staining (A). Type I fibers are green, type II fibers are black (unstained) and the dystrophin border is red. Scale bar is 500 µm. Fiber type composition (B), mixed fCSA (C), type I fCSA (D), type II fCSA (E), delta type I fCSA (F), and delta type II fCSA (G) pre- and post-intervention for HL-RET and LL-RET. Data are presented as means (bars) ± SD or with individual values (pre-post) from 12 subjects. Male subjects are presented as filled symbols (○), and female subjects are presented as unfilled (◯) symbols. Data was analyzed using two-way repeated measures ANOVA (B-E) and an unpaired test (F, G). ns; non-significant.

### Satellite cell and myonuclear content

Satellite cell content in mixed and type II fibers remained unchanged across the intervention (Figure 5A, C). However, in type I fibers, satellite cell content increased by ∼25% from pre-to post-intervention (main effect of time; p=0.014, figure 5B). In contrast, myonuclear content remained stable in type I fibers, but a slight decline was observed in mixed fibers (∼6%) and type II fibers (∼8%) from pre-to post-intervention (main effect of time; p<0.05), as shown in Figure 5D, F.

**Figure 5.**
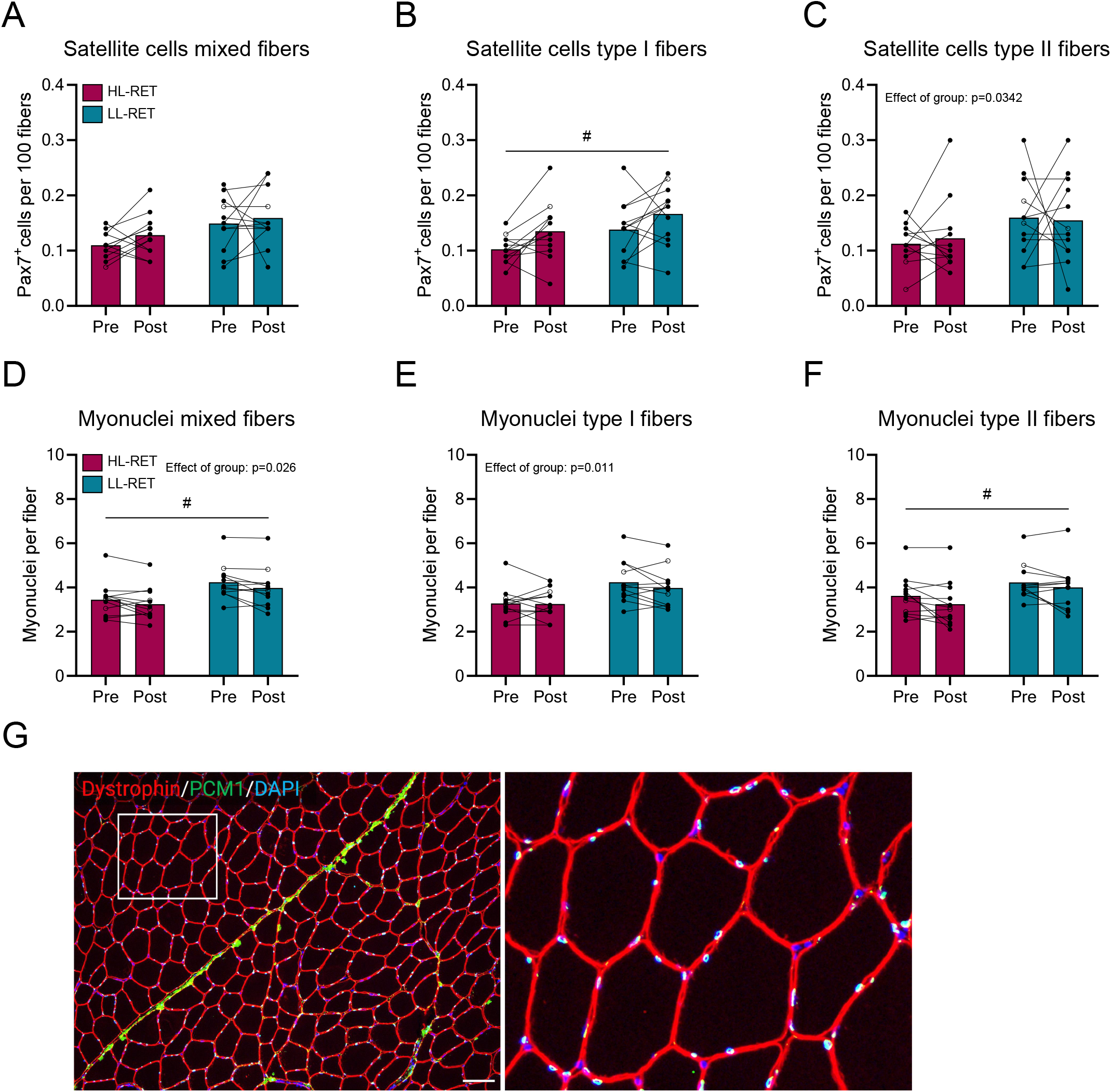
Satellite cell content in mixed fibers (A), type I fibers (B), and type II fibers (C) pre- and post-intervention for HL-RET and LL-RET. Myonuclear content in mixed fibers (D), type I fibers (E), and type II fibers (F) pre- and post-intervention for HL-RET and LL-RET. A representative image of the myonuclear staining (G). Dystrophin border in red, PCM1 in green and DAPI in blue. Scale bar is 100 µm. Data are presented as means (bars) with individual values (pre-post) from 12 subjects. Male subjects are presented as filled symbols (○), and female subjects are presented as unfilled (◯) symbols. Data was analyzed using two-way repeated measures ANOVA. A significant main effect of time is indicated by # p<0.05.

## Discussion

Muscular adaptations to LL-RET have been underexplored in trained individuals, with limited comparisons to traditional high-load training paradigms. This study aimed to investigate the muscular and myocellular adaptations of a 9-week LL-RET intervention (20-25 RM) and compare them to HL-RET (3-5 RM) in resistance-trained individuals, where all sets were taken to volitional failure. One of the key findings was that LL-RET, despite using significantly lower loads than traditionally recommended, produced strength gains comparable to HL-RET in multi-joint movements, but HL-RET led to superior strength gains in single-joint movements. The two distinct training regimens led to similar increases in muscle thickness, but at the myocellular level, neither type of training induced detectable muscle fiber hypertrophy. Only moderate changes were observed in the satellite cell pool, which expanded in a load-independent manner in type I fibers. Our data, therefore, suggests that LL-RET in trained individuals can lead to similar adaptations as a HL-RET regimen when it comes to maximum strength development and hypertrophic adaptations.

Manipulating training variables such as frequency, intensity, and volume is essential for optimizing adaptations to RET. However, because these variables are interdependent and influence one another, standardizing them across different training conditions can be challenging in scientific experiments. In the present study, we used an experimental approach where both conditions performed each set to volitional failure, as practiced also by other research groups (35, 38). This approach allowed the LL-RET condition to reach muscular failure across multiple sets while keeping the total number of working sets equal across conditions. However, as a result, the total training volume was not matched between the HL-RET and LL-RET conditions, with the latter condition lifting ∼1.5 to 3 times greater volume per session, depending on the exercise. In contrast, some studies comparing high-load and low-load training have used a volume-matched approach, where the low-load condition performs fewer working sets to match the total volume of the high-load condition (10, 22). However, this approach has consistently led to suboptimal adaptations in low-load conditions. For example, Burd and colleagues (2010) showed that low-loads were inferior for increasing rates of protein synthesis compared to high-loads when volume-matched, but not when the low-load condition performed a greater total training volume (9). Similar findings have been reported in chronic exercise studies, where volume-matched low-load training resulted in reduced muscle hypertrophy compared to high-load training (22). Considering these factors, we argue that the approach used in the current study offers a more precise comparison across conditions and accurately mirrors real-life training scenarios where multiple working sets are commonly employed.

Increased muscular strength is a fundamental outcome of RET and widely regarded as a key physiological adaptation, with benefits sought by individuals across a broad spectrum, from athletes to clinical populations. While improvements in strength are commonly associated with hypertrophic changes in muscle mass, they are also influenced by alterations in tissue architecture and neural adaptations (2). To optimally increase maximal strength in trained individuals, it is generally recommended that loads ≥70% of 1 RM are incorporated into the training regimen (2). In the present study, we observed substantial improvements in 1RM strength for both conditions in the leg press exercise (∼20% increase), while strength gains in the leg extension exercise were more pronounced in the HL-RET condition. These novel findings suggest that maximal strength, at least for multi-joint movements, can be increased to a similar extent with either 20-25 RM or 3-5 RM loads provided the exercises are performed to volitional failure. To our knowledge, this has not been previously demonstrated in trained populations, where most work support the use high loads for maximal strength development (48, 54). Nonetheless, some studies in trained men have indicated comparable strength gains regardless of loading strategy. For instance, Morton and colleagues reported that a whole-body LL-RET protocol (20-25 RM) performed over 12 weeks led to similar adaptations in 1 RM strength as a moderate-load regimen (8-12 RM) (38). It is important to note, however, that moderate-load regimens may not optimally stimulate 1RM strength development and thus could be considered suboptimal in this context. By contrast, our comparison of the LL-RET condition with a high-load protocol specifically designed to optimize maximal strength (HL-RET) allows us to conclude that training with 20-25 RM loads can lead to comparable, if not optimal, strength adaptations.

Interestingly, we did not observe parallel increases in 1RM strength for both conditions in the leg extension exercise. The reason for this discrepancy remains unclear though. On one hand, it may relate to the principle of task specificity, which perhaps is more applicable to single-joint movements where compensatory muscle actions are limited. Our data on local muscular endurance provides some support for this hypothesis. Specifically, at the end of the intervention, the LL-RET condition allowed for a greater number of repetitions performed at a submaximal load compared to the HL-RET condition, suggesting a relative advantage in terms of local muscular endurance by practicing with low loads. On the other hand, it is plausible that neural adaptations contribute more to multi-joint exercises and that lifting heavy loads would be more optimal compared to single-joint movements where the change in muscle force-producing capacity is the most deciding factor. Regardless of the underlying mechanism, these findings have practical relevance for coaches and practitioners seeking alternative strategies to optimize maximal strength in contexts where lifting heavy loads is not feasible, such as in clinical settings. The notion that similar adaptations can be achieved using diverse loading strategies has traditionally been associated with muscle hypertrophy alone. However, our results suggest that this principle may extend to strength gains in multi-joint exercises as well. Whether these distinct training regimens stimulate increases in muscular strength through the same physiological mechanisms remains an open question for future research.

Here we demonstrate that LL-RET induces robust hypertrophic adaptations over the course of a 9-week intervention as determined by increased muscle thickness (∼8%). This conclusion is further validated by muscle thickness measurements taken at two independent sites of the *vastus lateralis* muscle, which displayed cohesive results. Consistent with our initial hypothesis, muscle growth achieved through LL-RET is comparable to that seen with high-load training. This is particularly noteworthy given the distinct differences in repetition ranges between the two conditions (3-5 RM for HL-RET vs. 20-25 RM for LL-RET). These findings challenge the prevailing notion that muscle growth is primarily driven by moderate/high loads (2), suggesting instead that hypertrophy can be achieved across a broad spectrum of repetition ranges, provided the effort is high and sets are taken to volitional failure. As a result, the optimal repetition range for maximizing muscle hypertrophy is likely broader than previously recognized (2). This has significant implications for optimizing hypertrophic development in athletes and for communicating to the broader public that muscle growth is not solely dependent on the ability to lift heavy weights. This is especially relevant for individuals aiming to increase muscle mass but who cannot tolerate the mechanical stress associated with HL-RET, such as older adults experiencing joint discomfort or athletes recovering from injury. The finding that various training approaches—both high-and low-load—can produce comparable hypertrophic outcomes can be captured by the saying, “Many roads lead to Rome.”

Moreover, it is important to note that the potential for muscle growth through LL-RET has previously only been sparsely explored in the trained state (38, 48). This represents a critical area for investigation, as trained muscle tissue exhibit distinct acute molecular responses to RET compared to the untrained state and the potential for continued adaptation tends to diminish as training experience progress (36, 56). Additionally, previous research in subjects accustomed to RET has primarily focused on comparing low-load training to moderate-load conditions, typically within the 8-12 RM repetition range (38, 48). Our study thus expands the existing literature by demonstrating that skeletal muscle shows great flexibility for hypertrophic adaptation to various training stressors, even in the trained state. Our findings also raise new questions, such as whether incorporating LL-RET into a traditional training program could help alleviate stagnation among experienced lifters and potentially break through plateaus that often occur as training experience increases, as indicated previously by our research group using LL-RET with BFR (4). This represents an avenue for further research.

Here we evaluated changes in muscle mass by assessing muscle thickness using ultrasound and by measuring fiber size in muscle cross-sections. However, contrary to the observed changes in muscle thickness, no alterations in muscle fiber size were detected throughout the intervention, even when data from the two conditions were pooled (main effect of time for mixed fibers p=0.28). This discrepancy between various hypertrophic markers has been reported previously by several independent laboratories (21, 46). While the precise cause remains unclear, it is possible that the inherent limitations of muscle biopsy sampling contribute to this inconsistency. Specifically, it is assumed that a sample containing ∼250-1000 fibers represent changes occurring at the whole muscle level. In this regard, previous studies, including our own, have shown considerable variability in fCSA at different sites of the *vastus lateralis* muscle (24, 42, 52), even in the absence of an intervention, potentially explaining the lack of a hypertrophic effect in this study. Despite incorporating a large number of fibers (≥150 fibers per biopsy) in our fCSA analyses to counter this marked variability, we could not detect such a hypertrophic effect.

On the contrary, it is important to note that the discrepancy between the different measures in our study could, in theory, arise from ultrasound measurements potentially overestimating changes due to factors such as swelling or edema, as noted in previous research (13). However, given that the post-intervention scans were conducted ≥72 h after the final exercise session and that the participants were already accustomed to RET, we believe any impact from these factors was negligible.

The secondary objective of this study was to examine whether two distinct loading regimens elicit different hypertrophic responses in type I and type II muscle fibers. This investigation was to some extent motivated by a recent review, which concluded that there is insufficient evidence to support the assertion that different loading strategies preferentially induce hypertrophy in specific fiber types (19). However, since we were unable to demonstrate a clear hypertrophic effect at the muscle fiber level, we could not address this question conclusively. Previous studies in resistance-trained populations have shown that the growth of type I and type II fibers is equivalent, regardless of the loading strategy, i.e., low vs. moderate loads (38). This is likely due to the recruitment of the entire spectrum of fiber types as sets are taken to failure (18). Whether this holds true when contrasting more extreme loading regimens, such as low vs. high loads, although remains to be determined. Future research exploring this question in trained individuals may benefit from extended protocols, which could facilitate more pronounced hypertrophic adaptations at the myocellular level. Additionally, alternative methods for assessing fiber type-specific hypertrophy, beyond traditional analysis on muscle cross-sections, should be incorporated. Techniques such as assessments of muscle fiber volume in isolated single fibers could provide more comprehensive insights into these adaptations (23, 45).

Following an acute bout of resistance exercise, satellite cells are activated, proliferate, and expand their own cell pool (25). Over multiple sessions and depending on the stimuli, satellite cells can fuse with muscle fibers to donate their nuclei (28, 49). This process, leading to the accretion of myonuclei, is believed to support muscle growth and adaptation, though its precise role remains a subject of ongoing debate (40). Prior work has shown that LL-RET, when combined with BFR, induces rapid and pronounced increases in satellite cell number and myonuclear accretion (5, 14, 43). However, the effects of LL-RET alone, particularly in the trained state, remain less well understood (31, 37, 55). In this study, we observed that satellite cell pool expansion occurred in a fiber type-specific manner regardless of the training regimen. Specifically, satellite cells associated with type I fibers increased by ∼25%, while no significant effect was observed in type II associated satellite cells. These findings complement previous work suggesting that LL-RET leads to an acute expansion of the satellite cell pool (55). Our results also imply that satellite cell expansion is not dependent upon high-load training, as it occurs across various repetition ranges.

Moreover, our findings add to a growing body of evidence indicating that satellite cell pool expansion is uncoupled from changes in myofiber size (57), and myonuclear accretion (28). This supports the emerging concept of fusion-independent mechanisms of satellite cells, which have received attention in recent years (40). One possibility is that satellite cell pool expansion primes the muscle tissue for subsequent growth by facilitating intracellular signaling and contributing to the remodeling of the extracellular matrix (41), a key process for sustaining muscle fiber growth (7). These findings suggest that satellite cells play a broader role in muscle adaptability beyond their traditional role as precursors to new myonuclei. Finally, the observation that satellite cells expanded exclusively in type I fibers is interesting. This result may be attributed to the subjects’ prior experience with resistance exercise (minimum of 2 years), which likely recruited and put the type II fibers under significant stress for an extended time period (19). Consequently, the subjects in this study could have a greater adaptive potential left in their type I fibers. A similar phenomenon has been observed in trained powerlifters, where type I fibers exhibit greater responsiveness to training, potentially due to the myocellular adaptations of their fast-twitch fibers approaching a physiological ceiling (4).

In the present study, we observed no significant alterations in myonuclear number, if anything, a slight reduction was noted across the intervention. The absence of myonuclear accretion is likely attributed to the lack of a significant change in muscle fiber size. While this relationship can be variable, some studies have reported a positive correlation between these two variables (1, 44). From the perspective of the myonuclear domain hypothesis, which posits that additional myonuclei are recruited via satellite cells when muscle fiber size approaches a theoretical limit (28, 44), our findings suggest that the “demand” for myonuclear accretion was minimal in the current cohort. This is supported by the baseline myonuclear domains (∼1900 µm^2^), which fall below the threshold (≥2000 µm^2^) proposed to trigger the need for additional myonuclei to support myocellular growth (44). Thus, one might speculate that these individuals had already recruited an adequate number of myonuclei to sustain their existing myofiber size, thus requiring a more pronounced hypertrophic stimulus to reinitiate this process. However, the notion of rigid myonuclear domains has been challenged in recent literature, with current perspectives suggesting that the myonuclear domain theory is more flexible than initially proposed (39, 49).

In conclusion, when sets are performed to volitional failure, LL-RET and HL-RET lead to comparable increases in muscle thickness over a 9-week training period. In terms of functional performance, the LL-RET regimen leads to similar increases in maximal strength in multi-joint movements, but not in single-joint movements, and leads to superior increases in local muscle endurance, as compared to HL-RET. Additionally, both training protocols effectively expanded the satellite cell pool, particularly in type I fibers. Thus, our data suggest that LL-RET can replicate the effects of HL-RET in terms of muscle hypertrophic development and myocellular adaptations and may also serve as an alternative to traditional high-load programs for improving certain aspects of maximal strength. Our findings thus provide novel insights into the effects of LL-RET and HL-RET in resistance-trained individuals and highlights the notion that different training strategies may yield similar adaptations, like the saying “many roads lead to Rome.”

## Author contributions

N.P. and O.H. conceived and designed the research. R.K. and G.D. supervised the training intervention. K.T.C., I.C.E., R.K., and G.D. performed the experiments. T.R. and N.P. supervised the project. K.T.C., N.P., and O.H. analyzed the data. K.T.H., I.C.E., R.K., G.D., T.R., N.P., and O.H. interpreted the results of the experiments. O.H. prepared figures and drafted the manuscript. K.T.H., I.C.E., R.K., G.D., T.R., N.P., and O.H. edited and revised the manuscript, and all authors approved the final version of the manuscript.

## Acknowledgments

We acknowledge the participants for taking part in this study. Figure 1 was created with BioRender.com.

## Data availability

The data supporting the findings of the present study are available from the corresponding author upon reasonable request.

